# Functional limb muscle innervation prior to cholinergic transmitter specification during early metamorphosis in *Xenopus*

**DOI:** 10.1101/173328

**Authors:** F. M. Lambert, L. Cardoit, E. Courty, M. Bougerol, M. Thoby-Brisson, J. Simmers, H. Tostivint, D. Le Ray

## Abstract

In vertebrates, functional motoneurons are defined as differentiated neurons that are connected to a central premotor network and activate peripheral muscle using acetylcholine. Generally, motoneurons and muscles develop simultaneously during embryogenesis. However, during *Xenopus* metamorphosis, developing limb motoneurons must reach their target muscles through the already established larval cholinergic axial neuromuscular system. Here, we demonstrate that at metamorphosis onset, spinal neurons retrogradely labeled from the emerging hindlimbs initially express neither choline acetyltransferase nor vesicular acetylcholine transporter. Nevertheless, they are positive for the motoneuronal transcription factor Islet1/2 and exhibit intrinsic and axial locomotor-driven electrophysiological activity. Moreover, the early appendicular motoneurons activate developing limb muscles *via* nicotinic antagonist-resistant, glutamate antagonist-sensitive, neuromuscular synapses. Coincidently, the hindlimb muscles transiently express glutamate, but not nicotinic receptors. Subsequently, both pre- and postsynaptic neuromuscular partners switch definitively to typical cholinergic transmitter signaling. Thus, our results demonstrate a novel context-dependent re-specification of neurotransmitter phenotype during neuromuscular system development.

## INTRODUCTION

The features that define a specific neuronal phenotype are generally conserved between species, are specified early during development, but can also undergo adaptive plasticity related to activity after maturation (Demarque and Spitzer, 2012; Borodinsky et al., 2014). Amongst the large variety of neuronal phenotypes, the developmental properties of the motoneuronal class are particularly well established and are shared by all vertebrates. In mammals, motoneuron (MN) specification begins in the ventral neural tube with the induction of progenitors by sonic hedgehog proteins (Roelink et al., 1995), the graded concentration of which triggers the subsequent expression of specific post-mitotic transcription factors (Goulding, 1998; Jessell, 2000). The homeodomain-containing protein Islet1 is the first molecular marker of MN differentiation (Ericson et al., 1992) and induces the later expression of homeobox Hb9, which consolidates the motoneuronal phenotype and participates in MN migration and central motor column formation (Arber et al., 1999). Soon after their specification, MNs express the two typical proteins associated with cholinergic neurotransmission, choline-acetyltransferase (ChAT) and the vesicular acetylcholine transporter (VAChT), enabling them thereafter to activate their muscle targets (Phelps et al., 1991; Chen and Chiu, 1992). The muscles, which develop concomitantly and contribute to motoneuron axon path-finding, also play a fundamental role in MN phenotyping and survival (Yin and Oppenheim, 1992; Kablar and Belliveau, 2005). Finally, MNs are considered to be fully functional once they have become affiliated to a corresponding central motor network and provide impulse-elicited excitation to muscle fibers using acetylcholine (ACh) as the neurotransmitter (Davis-Dusenbery et al., 2014).

In vertebrates, axial MNs innervating trunk muscles are distributed rostro-caudally along the spinal cord in the medial motor column (MMC), whereas fore- and hind-limb MNs are located in the brachial and lumbar lateral motor columns (LMC), respectively. In the amphibian *Xenopus laevis*, the axial and appendicular MNs controlling tail and limb muscles respectively are generated during two separate developmental periods. The former develop during pre-hatchling embryonic stages and control larval undulatory tail-based swimming by projecting to and exciting axial myotomes via nicotinic ACh receptor activation (van Mier et al., 1985; Sharpe and Goldstone, 2000). Thus, axial neuromuscular ontogeny takes place under conditions equivalent to those in mammals where the MNs and target muscles of a primary motor system develop simultaneously. In contrast, MNs of the neuromuscular system responsible for later limb-based locomotion differentiate and the limb buds start developing during early metamorphosis (Marsh-Armstrong et al., 2004), when the axial MNs and tail muscles are already entirely developed and operational.

Given that the primary axial and secondary appendicular MNs of *Xenopus* initially share the same anatomical and neurochemical environment, and because such features influence neuromuscular development (Yang and Kunes, 2004; Menelaou et al., 2015), it is conceivable that the emergence of the secondary limb MNs and their target muscles is susceptible to influences exerted by the already fully established and functional axial neuromuscular system. We thus hypothesized that the axial myotomes and their innervation constitute a potential interfering environment for the newly developing appendicular neuromuscular system, and that consequently, the latter may have to follow particular developmental rules adapted to this unusual context. In the present study, therefore, we investigated the developmental strategy employed by the early developing limb MNs and associated neuromuscular junctions, with a specific interest in exploring their neurochemical phenotype and functional capability, both in terms of spinal locomotor circuit interactions and limb bud muscle control. Our results show that during a brief pre-metamorphic period, the emerging limb MNs transiently express a noncholinergic transmitter phenotype that involves glutamate, while exhibiting all other characteristics of typical vertebrate MNs.

## RESULTS

### Spatial organization of the appendicular spinal motor column during early metamorphosis

Appendicular MNs start to innervate the hindlimb buds as soon as the latter begin to emerge at stage 49. At stage 50, the nascent limb consists of a tissue protrusion that is visible on the ventral side of the rostral tail myotomes, next to the larva’s abdomen (Nieuwkoop and Faber, 1956). The limb bud then continues to grow and differentiate throughout the pre-metamorphic period, during which time the animal triples in size (Fig. 1A_1_). At stage 56/57, the adult-like hindlimb is formed with differentiated thigh and leg segments along with the appearance of five webbed toes (Fig. 1A_2_). The limb extensor and flexor muscle groups are also differentiated at this time.

**Figure 1.**
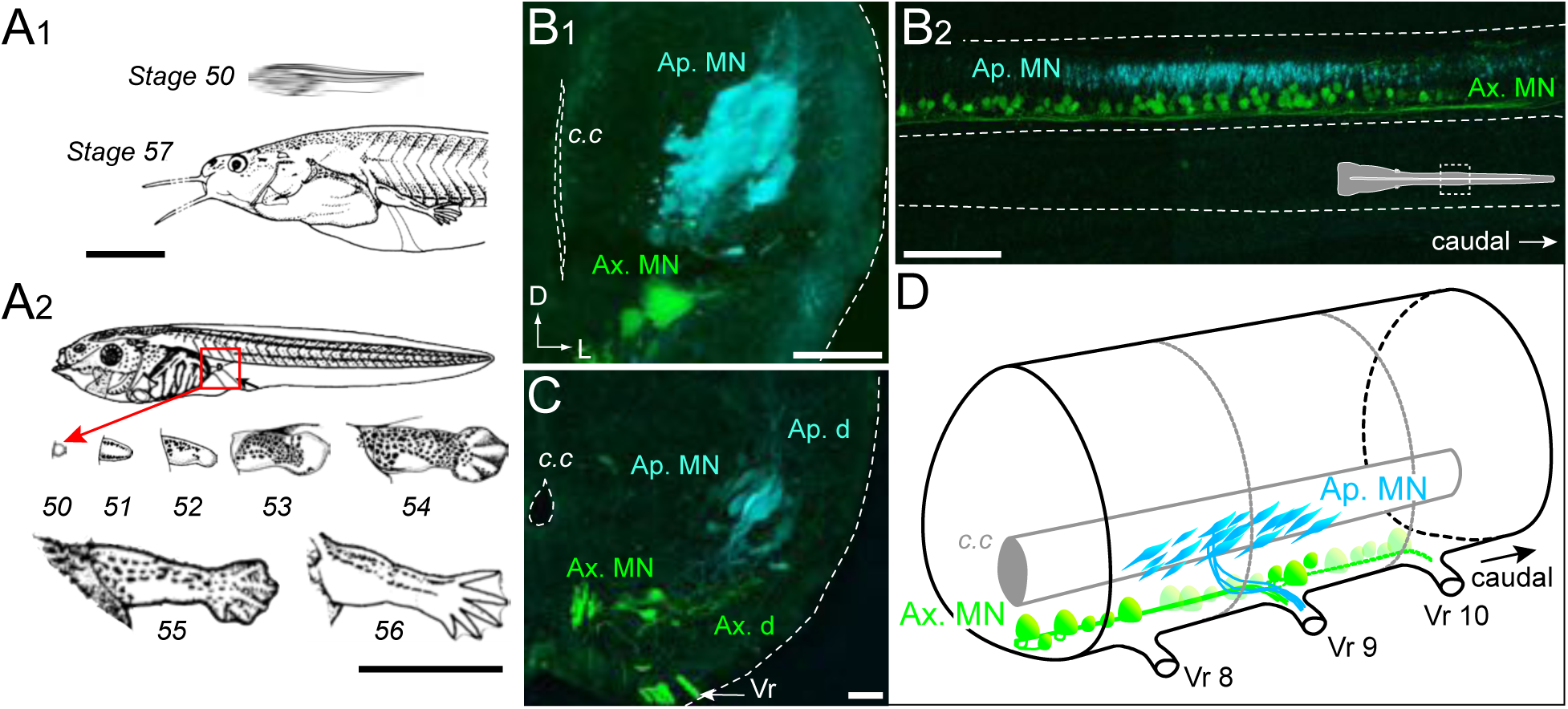
Developmental stages of *X. laevis* and anatomical organization of the spinal motor columns. A. Relative size differences between stage 50 and 57 larvae (A_1_) and associated morphological changes of the hindlimb bud during metamorphosis onset (A_2_). B, C. Segmental (B_1_, C) and rostrocaudal (B_2_) organization of the spinal motor column region containing limb MNs at stages 50-52 (B_1_, B_2_) and 53 (C). Inset in B_2_ shows the location in the spinal cord. D. Schematic representation of the segmental organization of the appendicular and axial motor columns in the larval spinal cord. Ap./Ax. MNs, appendicular/axial motoneurons; Ap./Ax. d., Ap./Ax. dendrites; Vr, ventral root; c.c, central canal; SC, spinal cord; D, dorsal; L, lateral. Scale bars: B_1_ and C = 20 μm; B_2_ = 100 μm.

To investigate the central spatial organization of the motoneurons innervating the hindlimb muscles during this developmental period, we injected two different retrograde tracers into a hindlimb bud and ipsilateral axial myotomes of stage 50 to 57 animals to label the somata and dendrites of appendicular and axial MNs, respectively. Each population is located centrally in separate motor columns. Specifically, the axial motor column is distributed ventro-medially along the entire length of the spinal cord, whereas the hindlimb motor column is located more medio-laterally and restricted to spinal segments 7 to 9, identified by counting caudally the number of ventral roots from the obex (Fig. 1B-D), as reported previously (Hulshof et al., 1987). In early stages 50-52 (Fig. 1B_1_, B_2_), retrograde labeling from the limb bud revealed a high density of MNs with relatively small cell bodies (<10 μm) and a reduced dendritic arbor, located more dorso-laterally than the axial MNs. From stage 53 onward (Fig. 1C), the appendicular motor column is positioned more laterally due to the enlargement of these spinal segments with a lower neuronal density. The appendicular MN somata then become bigger (20-50 μm) and acquire a characteristic elongated cell body shape and extended dendritic arbor, extending from a ventro-medial to dorso-lateral region of the hemicord (Fig. 1C).

### Delayed cholinergic transmitter phenotype expression in the developing appendicular motor column

The time course of ACh neurochemical ontogeny in appendicular MNs was first investigated using fluorescence immunochemistry for ChAT and VAChT expression in spinal cord cross-sections at different developmental stages (Fig. 2A, B). At stage 53 (Fig. 2A), appendicular MNs were not labeled at all with the ChAT and VAChT antibodies, whereas in the same spinal slices, axial MNs were strongly positive for the two proteins. In contrast, at later stages (*e.g*., stage 57 in Fig. 2B) both ChAT and VAChT were clearly expressed in appendicular MNs (see lower left insets) as well as in axial MNs. Overall, ChAT and VAChT were not immuno-detected in appendicular MNs until stage 55, in contrast to axial MNs which were immunoreactive throughout the early pre-metamorphic stages.

**Figure 2.**
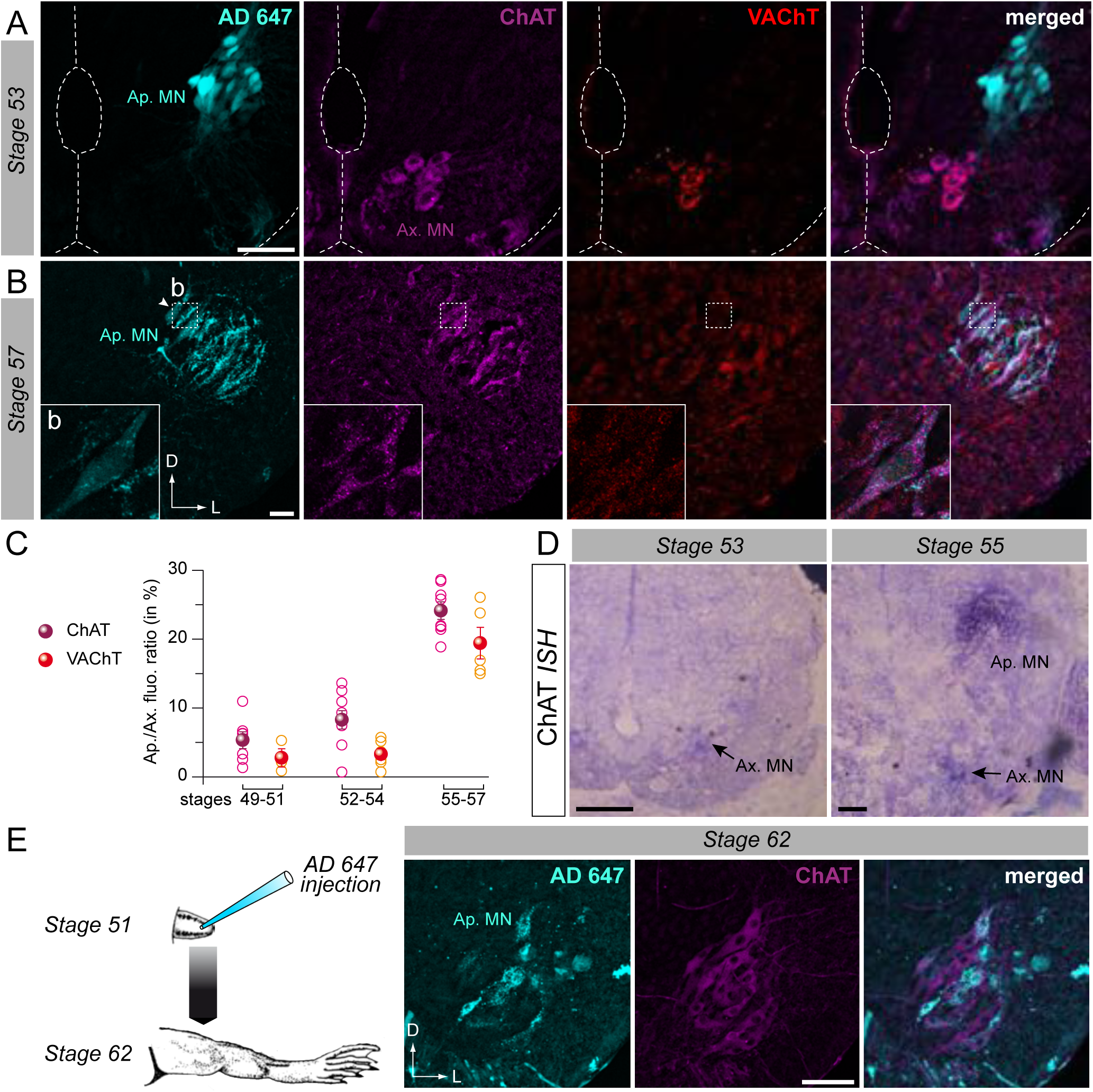
Developmental emergence of the molecular ACh phenotype in appendicular MNs. A, B. Examples of fluorescence immunolabeling against ChAT (magenta) and VAChT (red) in appendicular MNs (Ap. MN) labeled with retrograde Alexa Fluor dextran 647 (AD 647, cyan) in stages 53 (A) and 57 (B) tadpoles. Insets (b) in B show magnification of stage 57 appendicular MNs. C. Percentage ratio of fluorescence (fluo.) variation between appendicular (Ap.) and axial (Ax.) MNs for ChAT (magenta) and VAChT (red) immuno-signals from stages 49 to 57. Open and partially filled circles represent single preparation and group averages (± SE), respectively (see Methods). D. Examples of *in situ* hybridization (ISH) labeling for ChAT mRNA in the appendicular spinal column at stages 53 and 55. E. Example of fluorescence immunolabeling against ChAT in stage 62 appendicular MNs (Ap. MN) which were previously labeled with AD 647 injected into the hindlimb at stage 51 (see schematic at left). All scale bars = 50μm; D, dorsal; L, lateral.

Semi-quantitative analysis of ChAT/VAChT fluorescence intensity in axial and appendicular MNs (see Methods) was next performed on confocal image stacks in which appendicular MNs were identified by retrograde tracer labeling. The appendicular/axial fluorescence ratio measured at stages 49-54 showed that the immuno-signal detected in the appendicular motor column was not significantly higher than the background signal level (ChAT: 5.3 ± 1.2 at stage 49-51; 8.3 ± 1.3 at stage 52-54; VAChT: 2.8 ± 1.3 at stage 49-51; 3.3 ± 0.8 at stage 52-54; Fig. 2C). Immuno-labeling in the LMC became more robust with further larval development, with the appendicular/axial fluorescence ratio increasing to ~20% (at stage 55-57; ChAT: 24.1 ± 1.3; VAChT: 19.4 ± 2.3; Fig. 2C), which was consistent with the clear detection of both ChAT and VAChT in appendicular MNs at the later pre-metamorphic stages.

Because *in situ* hybridization (ISH) is more sensitive than immuno-detection, ISH was also performed to detect ChAT mRNA in a group of larvae at stages ranging from 49 to 57. As illustrated in Fig. 2D, no signal was detected in the appendicular motor column area up to stage 53, whereas a ChAT ISH signal became clearly evident from stage 55 onwards. In contrast, a strong ChAT ISH signal was observed in axial MNs over the entire developmental range examined. Thus both these immunochemistry and ISH results confirmed that appendicular MNs do not exhibit the cholinergic molecular phenotype prior to stage 55, although they innervate the hindlimb bud muscles from stage 49.

One possibility was that the non-cholinergic and cholinergic appendicular MNs observed at different developmental stages were in fact distinct populations. To address this possibility we first applied a fluorescent retrograde dye to the hindlimb bud at stage 51/52, prior to the appearance of the ACh phenotype in limb MNs (Fig. 2E). Thereafter, the tracer-treated larvae were raised until they reached metamorphosis climax, by which time the appendicular system is fully developed. ChAT immunolabeling then performed on spinal cross sections taken from these stage 62 animals strongly labeled the entire appendicular motor population (Fig. 2E, middle panel). Significantly, some of these ChAT-positive MNs were also co-labeled with the retrograde fluorescent dye previously applied at earlier stage 51/52 (Fig. 2E, right panel). Thus, these double-labeled MNs that innervated the hindlimb at stage 62 were already present at stage 52, before they expressed the cholinergic molecular phenotype. This finding therefore demonstrated that early non-ACh appendicular MNs constitute at least a sub-population of the cholinergic MNs that constitute the future mature appendicular motor innervation after metamorphosis.

### Molecular and functional identification of early non-cholinergic appendicular neurons

The molecular identity of the non-ACh neurons innervating the newborn limbs at early pre-metamorphic stages was verified by immunochemistry against the transcription factor Islet1/2, a specific protein marker of motor neurons (Ericson et al., 1992). Double immunostaining in larvae younger than stage 55 revealed that the ChAT-negative neurons labeled with retrograde dye applied in the hindlimb bud were also strongly Islet1/2-positive, similar to the mature axial MNs present in the same slices (Fig. 3A).

**Figure 3.**
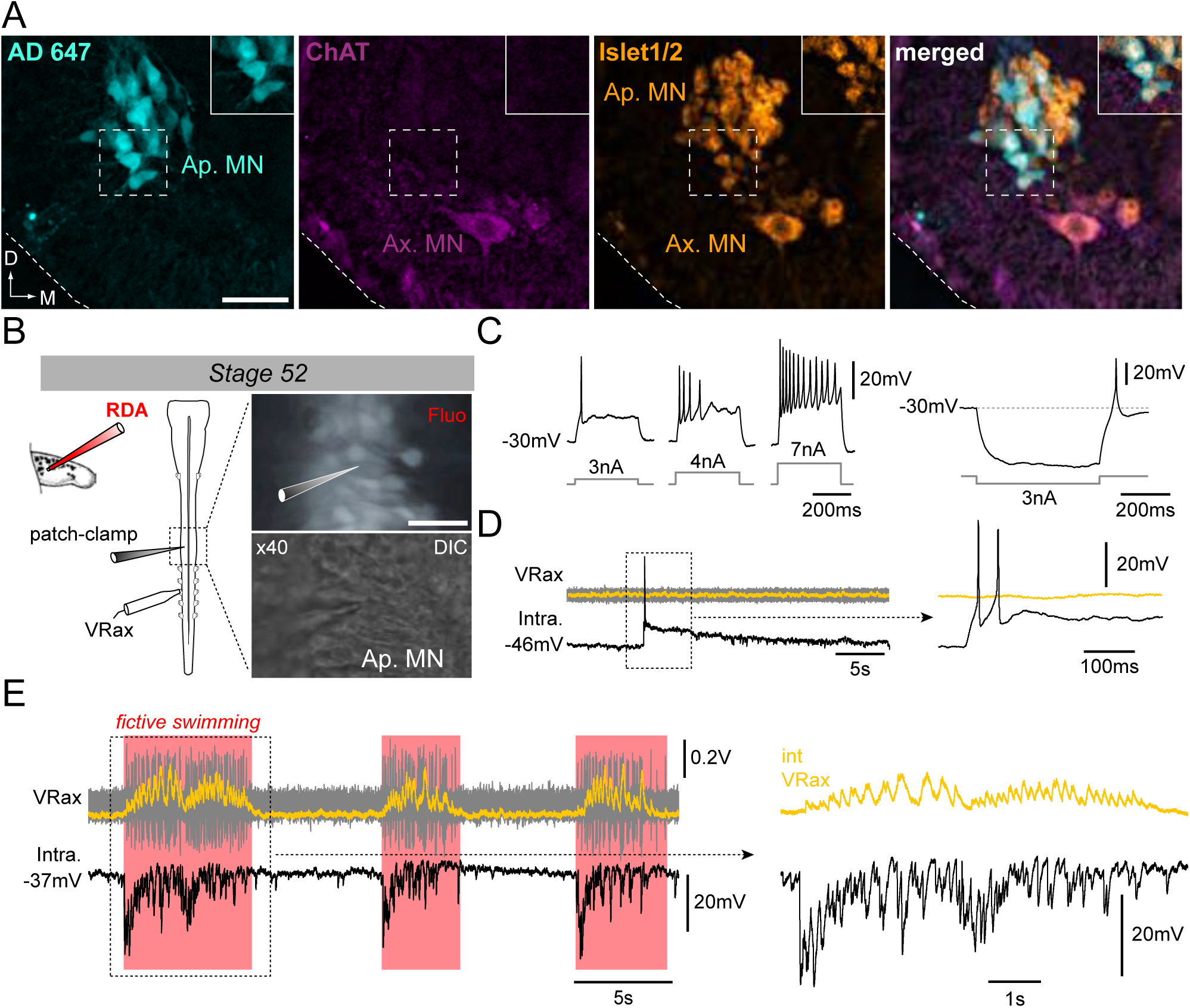
Motoneuronal identification of non-cholinergic limb projecting neurons. A. Example of fluorescence immunolabeling against ChAT (magenta) and Islet1/2 (orange) in appendicular and axial MNs (Ap. MN, Ax. MN) in a stage 52 tadpole. Appendicular MNs were previously labeled with retrograde Alexa Fluor dextran 647 (AD 647, cyan). B. Intracellular recordings from appendicular MNs. Stage 52 MNs were identified with prior rhodamine dextran amine (RDA) retrograde labeling from the hindlimb bud and recorded with cell-attached patch-clamp in whole CNS preparations in order to preserve spinal locomotor circuitry. C. Increasing steps of injected depolarizing current elicited increasing spike discharge while hyperpolarizing current injection evoked rebound spiking. The MN (Intra) could also fire spontaneously in the absence of axial ventral root (VRax) activity (D), and received strong rhythmic synaptic input during spontaneous episodes of fictive axial locomotion (E, see red shading). A single episode on an extended time scale is shown at right. VRax traces (in yellow) are integrated transforms of raw extracellular VR recordings. All scale bars = 50μm; D, dorsal; M, medial.

The electrophysiological properties of these non-ACh appendicular MNs were then tested at stage 52 by performing patch-clamp intracellular recordings of neurons identified by RDA retrograde labeling from the hindlimb bud (Fig. 3B). These recorded MNs (n=11) had an average resting membrane potential of 40.2 ±8.2 mV and were able to produce action potentials either in response to increasing steps of depolarizing current injection (Fig. 3C, left), on rebound after release from experimental hyperpolarization (Fig. 3C, right) or even spontaneously (Fig. 3D). Significantly, moreover, during spontaneous repetitive episodes of locomotor-like activity monitored from a more caudal spinal ventral root, correlated synaptic fluctuations were observed in all identified appendicular MNs, thereby indicating a functional synaptic coupling with other components of the spinal locomotor network (Fig. 3E).

Altogether, these findings demonstrate that the retrogradely-labeled, non-ACh spinal neurons that prematurely innervate hindlimb buds express the motoneuronal marker Islet1/2 and present basic biophysical characteristics of functional MNs, being able to produce impulses as a function of membrane potential that in turn can be influenced via synaptic influences from central premotor circuitry.

### Locomotor-related activation of hindlimb muscles by non-cholinergic motoneurons

Whether the early appendicular MNs are actually capable of driving muscle activation in the hindlimb buds before the emergence of the cholinergic phenotype was next investigated by making EMG recordings from both limb bud muscles and axial myotomes in semi-intact larval preparations (Fig. 4A left panel). During episodes of spontaneous axial fictive swimming in a stage 53 preparation (Fig. 4A, right panel), locomotor commands monitored from a caudal spinal ventral root elicited rhythmic EMG activity in both the segmental myotomes and limb bud muscles. The ventral root bursts occurred in phase with EMG bursts in the recorded ipsilateral hindlimb muscle and in phase opposition with bursts in the contralateral myotome (Fig. 4B, control). Under subsequent bath application of *d*-tubocurarine to block any nicotinic receptors and thereby prevent ACh-dependent synaptic transmission (Sillar and Roberts, 1992, 1993), the expression of swimming-related ventral root burst activity persisted, but associated tail myotome EMG activity was completely abolished after 15 min of antagonist perfusion (Fig. 4B, *d*-tubocurarine). In contrast, even after >30 min *d*-tubocurarine perfusion the hindlimb bud muscles continued to exhibit rhythmic EMG discharges that, although reduced in amplitude, remained strictly coordinated with the ongoing axial pattern (Fig. 4B, *d*-tubocurarine; see black arrow in lower middle plot). The ability of both the tail myotomes and limb bud muscles to express locomotor-related activity recovered fully after 2 h washout with normal saline (Fig. 4B, wash). Semi-intact preparations older than stage 54 similarly exhibited EMG burst activity in their hindlimb buds and tail myotomes during fictive axial locomotion in control conditions (e.g., stage 56 in Fig. 4C, left). In these cases, however, a 15 min bath application of *d*-tubocurarine decreased the frequency of the centrally-generated axial swim pattern, and completely abolished, albeit reversibly, associated EMG activity in both the hindlimb bud muscles and tail myotomes (Fig. 4C, *d*-tubocurarine; wash).

**Figure 4.**
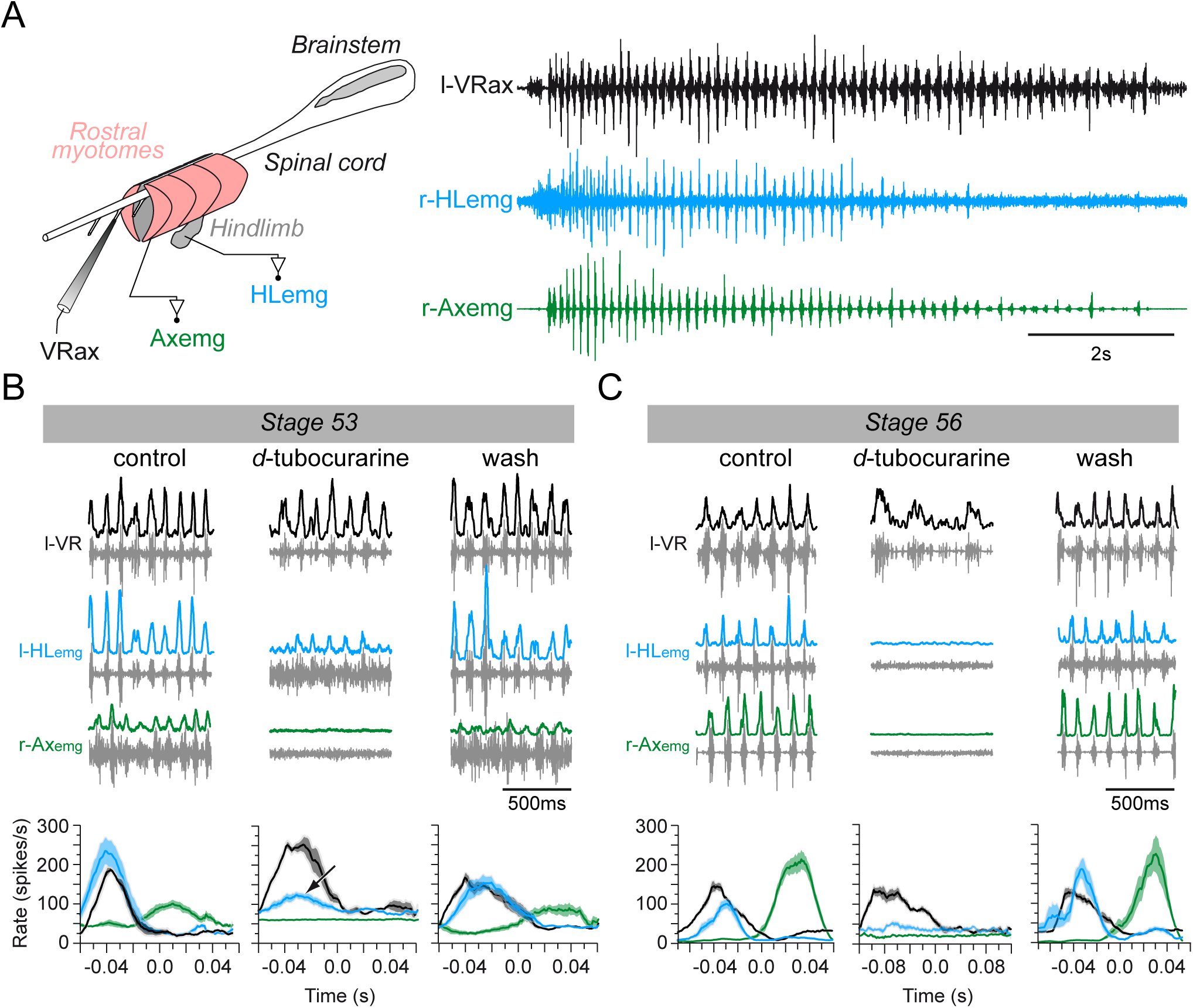
Switch from non-cholinergic to cholinergic limb muscle activation during axial swimming at different developmental stages. A. Rhythmic burst discharge (right panel) recorded from a left side axial ventral root (l-VRax) together with a right side hindlimb bud muscle (r-HLemg) and axial myotome (r-Axemg) during a spontaneous Active swim episode in a semi-isolated stage 53 preparation (left panel). B, C. Examples of recordings before (control), during and after (wash) bath application of *d*-tubocurarine to a stage 53 (B) and a stage 56 (C) preparation. Lower plots in B and C show instantaneous l-HLemg (blue) and r-Axemg (green) discharge rates (spikes/s) averaged (± SE) over 10-20 l-VRax locomotor cycles (black). Black arrow in the middle graph of B indicates persistent limb bud EMG activity occurring in phase with the ipsilateral ventral root during *d*-tubocurarine application, and which was no longer present at the later developmental stage (c.f., middle plot in C).

The differential effects of *d*-tubocurarine at early and later larval stages thus indicated that a developmental shift in the mechanism by which the appendicular MNs activate their target muscles during fictive locomotion occurs around intervening stages 54-55. Whereas neuromuscular transmission is initially largely independent of ACh signaling in the pre-metamorphic tadpole, it changes to a completely ACh-dependent process in older tadpoles. This in turn suggests that developmental modifications in parallel with the switch in MN transmitter signaling must also occur in the receptor phenotype of the hindlimb muscles themselves.

### Developmental switch in hindlimb neuromuscular transmission

In the *Xenopus* embryo, immature axial MNs have been shown to co-express ACh and glutamate, although glutamate is reported to have no postsynaptic bioelectrical effects (Fu et al., 1998). Since our present data show that the appendicular MNs are able to activate hindlimb muscles before using ACh as a neurotransmitter, we asked whether glutamate may be involved in the early neuromuscular junction (NMJ) transmission process.

First, we explored in whole-mount hindlimb buds of various developmental stages (from 51 to 57) the putative expression and relative distribution of nicotinic and glutamate receptors with fluorescent α-bungarotoxin and the anti-NR2b antibodies, respectively (Fig. 5A). At stage 52, there was a total lack of labeling for α-bungarotoxin in hindlimb buds (Fig. 5A, upper panels), indicating the absence of muscle ACh receptors at this stage. In contrast, at the same developmental stage, diffuse NR2b fluorescence was observed in the vicinity of MN axons (visualized with neurofilament immuno-detection). It is noteworthy that a similar expression of NR1, another glutamate receptor subunit, was found which also matched parallel synaptophysin labeling (Fig. 5B). Thus, at this early pre-metamorphic stage, glutamate receptors were found to be present on hindlimb muscle fibers, close to synaptophysin-marked presynaptic terminals, suggesting the existence of functional glutamatergic NMJs.

**Figure 5.**
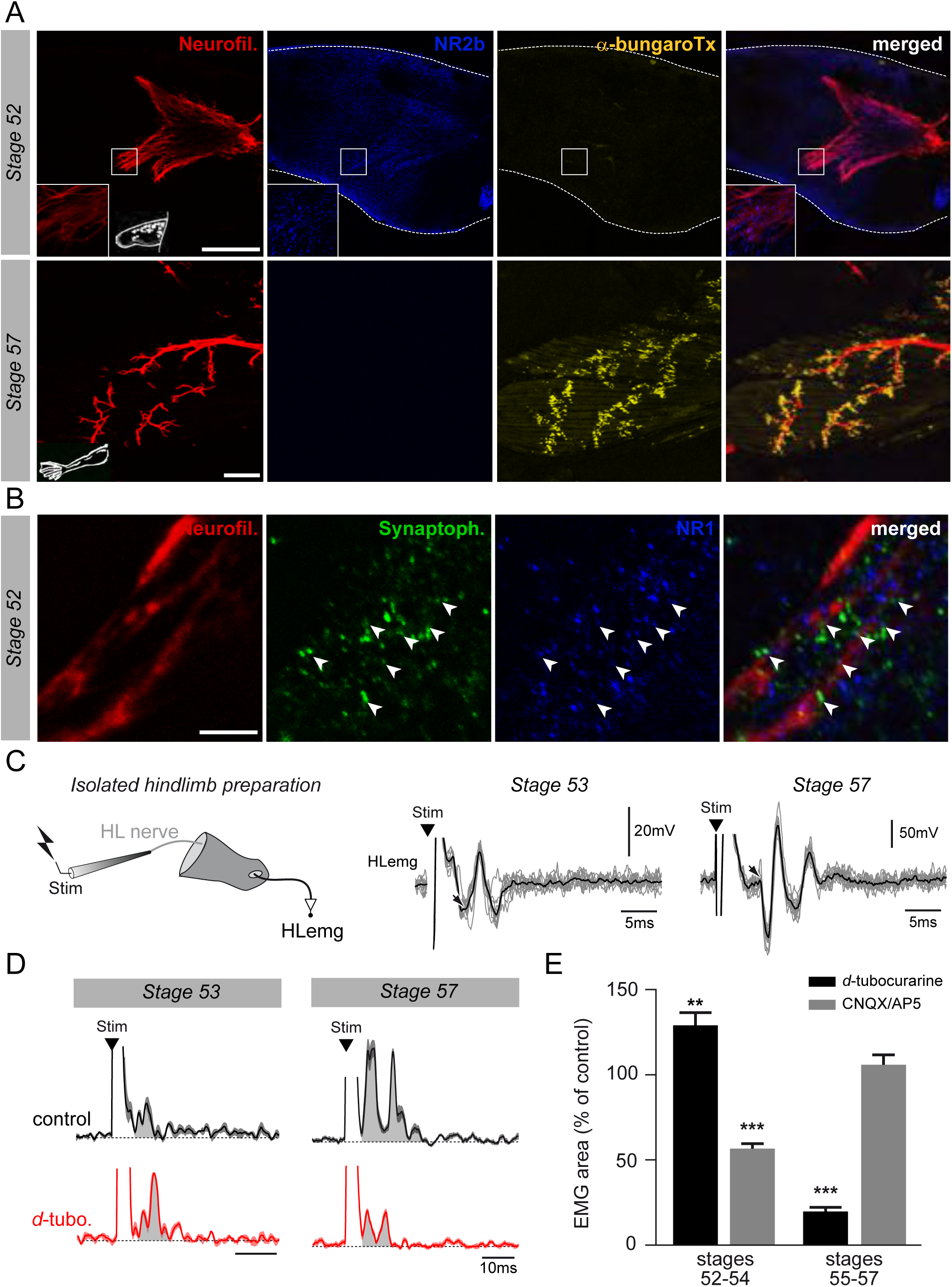
Molecular and functional switch in hindlimb neuromuscular transmission. A. Examples of hindlimb innervation patterns and distribution of ACh nicotinic receptors in a whole-mount hindlimb bud, revealed by fluorescence immunolabeling of neurofilament associated protein (Neurofil., red), glutamate receptor (NR2b, blue) and α-bungarotoxin labeling (α-bungaroTx, yellow) respectively, at stages 52 (upper panels) and 57 (lower panels). Inset drawings at bottom left of each panel show bud morphology at the two representative larval stages. Scale bars: 200μm for stage 52, 100μm for stage 57. B. Examples of fluorescence immunolabeling against neurofilament associated protein (Neurofil., red), synaptophysin (Synaptoph., green) and NMDA glutamate receptor subunit 1 (NR1, blue) in whole-mount limb bud at stage 52. Scale bar: 50μm. C. EMG recordings (HLemg) from an isolated hindlimb preparation (schematic at left) in response to single pulse (100μs) electrical stimulation (Stim) of a limb motor nerve (HL nerve) in stages 53 and 57 larvae. In each case, the black arrow indicates the beginning of the EMG response and the black trace represents the mean profile of 6 superimposed responses. D. Integrated motor nerve-evoked HLemg responses before (control) and during *d*-tubocurarine (*d*-tubo.) bath application to stage 53 and 57 preparations. Black traces represent the mean response profile (±SE); the area under each curve (grey) was used to measure the response size. E. Mean EMG response area as percentage of control response during *d*-tubocurarine (black) or CNQX/AP5 (grey) bath application at stages 52-54 and 55-57, respectively. ^∗∗^ p<0.01 and ^∗∗∗^ p<0.001, Mann–Whitney U-test.

The first α-bungarotoxin positive signal was detected at stage 55 with characteristic dispersed dots (data not shown), as previously described (Marsh-Armstrong et al., 2004). By stage 57, typical clusters of nicotinic receptors were observed close to appendicular nerve terminal branches, while NR2b (Fig. 5A lower panels) and NR1 receptor subunits (not illustrated) were no longer detected in the hindlimb bud. These findings therefore indicate that glutamate receptor subunits are no longer expressed from stage 57 onwards, but rather, are superseded by nicotinic receptors in likely correspondence to the switch in NMJ transmission to a cholinergic phenotype.

In a final step, the electrophysiological response properties of neuromuscular transmission were tested on isolated hindlimb preparations over a similar range of developmental stages (from 52 to 57; Fig. 5C). Single pulse electrical stimulation applied to a limb bud motor nerve elicited EMG responses in the hindlimb bud with a mean latency of 9.3 ± 2.6 ms at stages 52-54 and 5.1 ± 2.1 ms at stages 55-57, and a mean amplitude of 0.30 ± 0.06 mV at stages 52-54 and 2.49 ± 0.99 mV at stages 55-57. Either *d*-tubocurarine or CNQX/AP5 cocktail was then bath-applied to test whether these neuromuscular responses were ACh- or glutamate-mediated, respectively. At stages 52-54, the appendicular nerve stimulus-evoked EMG response was not diminished by subsequent *d*-tubocurarine application, but rather, was increased by 29 ± 8% (Fig. 5D, left; 5E, left black bar). [Note that such a *d*-tubocurarine-induced enhancement of post-junctional responses has already been reported in other animal models and is unrelated to its antagonistic effects on nicotinic receptors themselves (Egan et al., 1993; Baron et al., 1996)]. Conversely, the hindlimb muscle response was decreased by 43 ± 3% following bath-application of CNQX/AP5 (Fig. 5E, left grey bar). At stages 55-57 on the other hand, the nerve stimulus-evoked EMG response was decreased by 80 ± 2% under *d*-tubocurarine perfusion but was not significantly affected (p = 0.5) in the presence of CNQX/AP5 (Fig. 5D, right; 5E, right black and grey bars, respectively).

Altogether, these immunochemical and pharmacological results showed that glutamate, but not ACh, is initially involved in hindlimb muscle activation, whereas NMJ transmission becomes ACh-mediated and increasingly efficient (increase in response amplitude, reduction in transmission delay) from stage 55 onwards. This therefore confirms the occurrence of a functional switch in *Xenopus* hindlimb muscle properties that is commensurate with the switch in MN neurotransmitter phenotype during the pre-climax phase of metamorphosis.

## DISCUSSION

In this study we report that as early as pre-metamorphosis (i.e., before stage 55), *de novo* appendicular MNs are immunoreactive for MN-identifying Islet1/2 transcription factor, exhibit characteristic MN electrical properties and synaptic influences, are involved in locomotor bouts of activity, and project to limb bud muscles to evoke typical post-junctional responses. Unexpectedly, however, these MNs do not at this premature stage express the standard cholinergic neurotransmitter phenotype, but rather, they use glutamatergic neurotransmission to activate their target muscles. From between stages 55 and 57, however, the limb MNs and their NMJs simultaneously undergo a switch to acetylcholine-dependent signaling. Thus our data show for the first time that in metamorphosing *Xenopus*, MNs that innervate the newly emerging hindlimbs first develop through the employment of a transient, but functional, glutamatergic transmitter mechanism before maturing into a more conventional and definitive cholinergic phenotype.

Classically in vertebrates, developing MNs are perpetually cholinergic and extend their axons out of the spinal cord towards target muscles to form functional NMJs (Phelps et al., 1991; Goulding, 1998). However, in *Xenopus* the ontogeny of appendicular MNs does not obey this well-established developmental pattern, since the present data indicate that before the acquisition of a cholinergic phenotype, these MNs are able to transmit centrally-generated motor commands to the developing limb bud muscles using glutamate as the transmitter effector. However, this situation requires the concomitant expression of appropriate receptors at the post-junctional level in correspondence with the type of pre-synaptic neurotransmitter released. Accordingly, we observed that the use of MN glutamate prior to ACh is also matched by the presence of glutamate receptors prior to the appearance of cholinergic receptors at the limb NMJs. Such a parallel developmental sequence is therefore indicative of a complete neurotransmitter phenotype switching, whereby intrinsic molecular processes lead to a change of a transmitter to another within the same presynaptic neuron and a concomitant modification in associated postsynaptic receptor subtype in order to ensure synapse functionality. This loss/gain in pre-synaptic neurotransmitter/post-synaptic receptor phenotype has been reported to occur in developmental, post-lesional or activity-dependent contexts and is considered to be a major underlying feature of neuronal plasticity (Spitzer, 2017). It can either maintain or invert synaptic sign (Borodinsky et al., 2004; Borodinsky and Spitzer, 2007) and is generally associated with observable behavioral changes (Sillar et al., 1998; Demarque and Spitzer, 2010).

In *Xenopus*, limb MNs are born during pre-metamorphosis, then their number diminishes after the establishment of NMJs (Hughes, 1961; Prestige, 1967). This in turn raises the possibility that the initial non-cholinergic motoneuron population we identify here constitutes a short-lived subclass of LMC neurons that project transiently into the limb buds at pre-metamorphic stages, but then is replaced by a different cholinergic population as metamorphosis proceeds. However, our finding that limb bud MNs previously retrogradely-labeled at stage 51 are still present at stage 62 when the population of appendicular MNs is fully established and uniformly cholinergic (Baldwin et al., 1988), including our early-labeled neurons, argues against this possibility. The possibility that the initial non-cholinergic phenotype may be restricted to a specialized motoneuronal sub-population that innervates limb muscle spindles is also unlikely: frog spindles are mainly innervated by collateral branches of skeletal MNs (Katz, 1949; Gray, 1957) and although a possible specific fusimotor innervation has been proposed (Fujitsuka et al., 1987), it would constitute a very distinct population, the small proportion of which does not correspond to the large number of early non-cholinergic LMC neurons that we found. Thus, our data strongly indicate that the same motoneuronal population in the developing appendicular motor system of *Xenopus* does indeed utilize consecutively two different neurotransmitter signaling mechanisms in association with the emergence of hindlimb motility (Hughes and Prestige, 1967) and forthcoming limb-based locomotor and postural control (Combes et al., 2004; Beyeler et al., 2008).

Given that limb MNs acquire their cholinergic phenotype around stage 55 and remain cholinergic throughout adulthood (Baldwin et al., 1988), the existence of an early, transient but functional non-cholinergic phenotype is at first sight puzzling. In a broader context, a number of studies have reported the co-existence of various neurotransmitters in vertebrate MNs. For instance, axial MNs of *Xenopus* embryos co-express glutamate and ACh (Fu et al., 1998), while developing myotome fibers initially express a variety of receptors, including both cholinergic and glutamate subtypes, until NMJ formation when solely cholinergic receptors are preserved (Borodinsky and Spitzer, 2007). Co-released glutamate regulates the development and function of cholinergic neuromuscular synapses in larval zebrafish (Todd et al., 2004) and *Xenopus* (Fu et al., 1998) by potentiating ACh release through an activation of presynaptic receptors on the MN terminals themselves. Moreover, mammalian spinal MNs co-release glutamate and ACh centrally to activate Renshaw cells (Mentis et al., 2005; Nishimaru et al., 2005; Lamotte d’Incamps and Ascher, 2008), as well as at their peripheral terminals (Waerhaug and Ottersen, 1993; Rinholm et al., 2007) where post-synaptic muscle fibers possess both ACh and glutamate receptors (Mays et al., 2009). Here again, however, in no such case has glutamate been reported to produce muscle activation *per se*, but rather, this transmitter acts indirectly by regulating ACh’s own impact at the NMJ (Vyas and Bradford, 1987; Malomouzh et al., 2003; Pinard et al., 2003) *via* the activation of post-synaptic NMDA receptors (Pinard and Robitaille, 2008; Petrov et al., 2013). On the other hand, glutamate is the predominant excitatory neurotransmitter in the vertebrate CNS, and supraspinal glutamatergic neurons can re-specify functional glutamatergic NMJs from otherwise purely cholinergic synapses on mammalian skeletal muscles following grafting with transected peripheral motor nerve (Brunelli et al., 2005). Given such a short-term reorganizing capability and the fact that glutamate is a major excitatory neurotransmitter at the NMJ of phylogenetically distant invertebrates (Gerschenfeld, 1973), it is possible that the transient expression of glutamatergic neuromuscular transmission in pre-metamorphosing *Xenopus* is representative of a latent ancestral step in the evolutionary transition of intrinsic molecular programming of the NMJ to cholinergic-dependent signaling.

A further and more appealing possibility is that the switching process is related to early appendicular MN axon path-finding and initial NMJ formation because of the unusual context of frog metamorphic development where secondary limb MNs must migrate to their muscle targets with the primary axial neuromuscular apparatus already in place, fully functional and using ACh as the NMJ transmitter (Fig. 6). Transplantation experiments in *Xenopus* have demonstrated the ability of MNs to innervate novel tissue targets (Elliott et al., 2013). Moreover, amongst the many guidance cues that orient axon elongation during development, both target-derived signals and axon growth cone-released ACh participate in target reaching and initial synapse formation (Yin and Oppenheim, 1992; Erskine and McCaig, 1995; Yang and Kunes, 2004). On this basis, therefore, it is conceivable that the pre-existing axial neuromuscular system in *Xenopus* provides a potential disturbing environment that could attract appendicular motor axons, if they were cholinergic, to make improper neuromuscular connections during pre-metamorphic development. On the contrary, a matched expression of glutamatergic neurotransmitter and receptors in appendicular MNs and limb muscles during pre-metamorphosis could ensure the correct orientation of the developing axon growth cones towards the limb buds and the establishment of synaptic contacts appropriate for the control of limb movements (Fig. 6A). Supporting this hypothesis are previous findings that morphologically normal neuromuscular connections can still develop despite the experimental suppression of either pre- or postsynaptic cholinergic partners (Westerfield et al., 1990; Misgeld et al., 2002) and that glutamatergic CNS neurons can replace cholinergic MNs and reinnervate skeletal muscle fibers after a peripheral nerve lesion (Brunelli et al., 2005). In *Xenopus*, once the initial appendicular NMJs are established, both pre- and postsynaptic partners could thereafter be instructed to switch to their definitive cholinergic phenotype (Fig. 6B).

**Figure 6.**
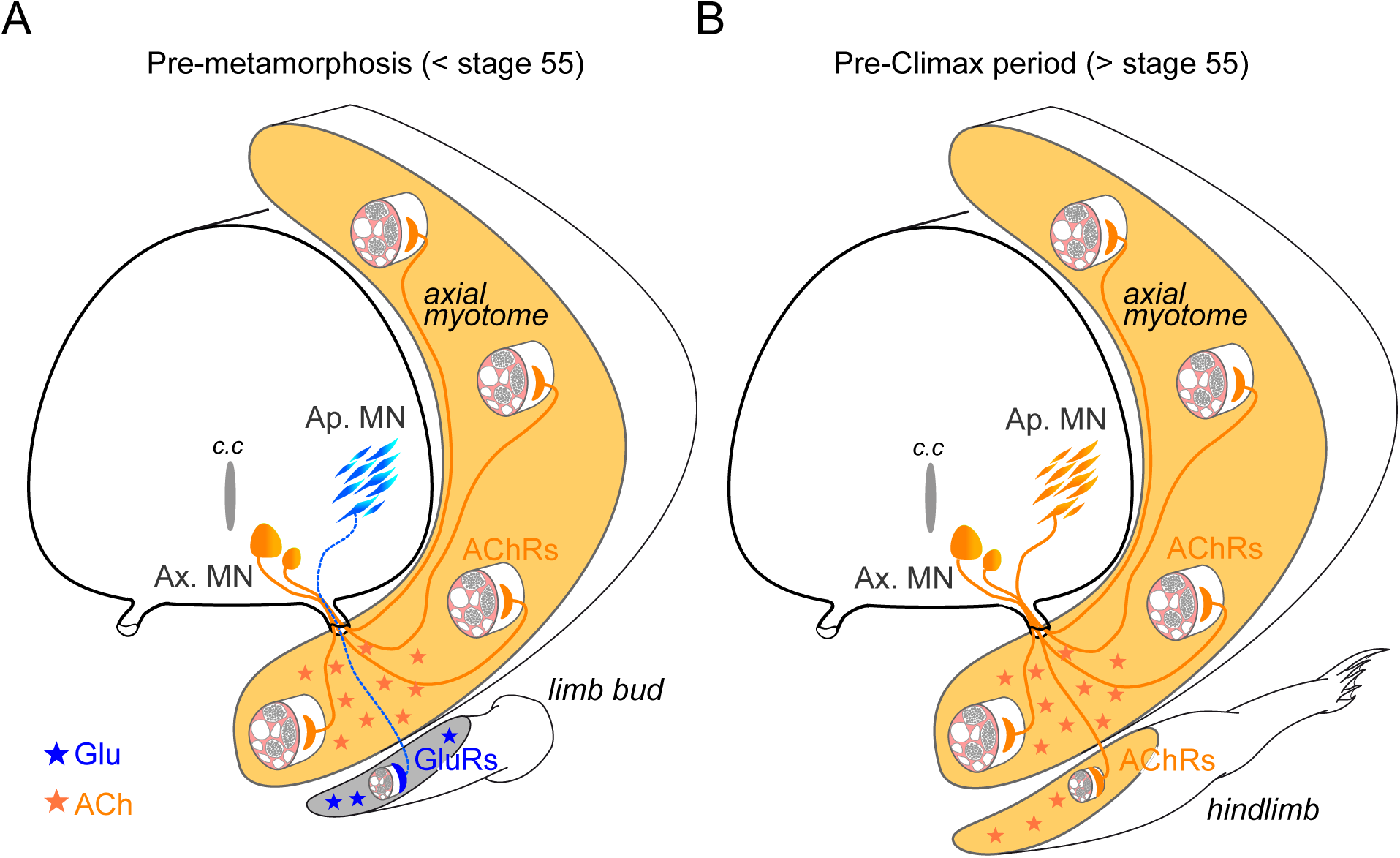
Schematic representation of neurotransmitter phenotype switching associated with the establishment of functional limb muscle innervation at different stages of *Xenopus* metamorphic development. See text for further explanation. AChRs: nicotinic ACh receptors; GluRs: glutamate receptors; Ap. MN: appendicular MNs; Ax. MN: axial MNs; c.c: central canal

The signal for such neurotransmitter re-specification remains to be determined, but thyroid hormones (TH), which control the developmental expression of ChAT (Patel et al., 1987; Gould and Butcher, 1989) and probably also VAChT since both proteins share the same gene locus (Eiden, 1998), are most likely to be involved. Consistent with this possibility is that the increase in thyroid hormone levels at metamorphosis onset, which starts at stage 54-55 in *Xenopus* (Shi, 2000), triggers a variety of gene-switching molecular programs required for the development of limb muscles and associated motor circuitry through the regulation of spinal cord neurogenesis and functions (Das et al., 2002; Marsh-Armstrong et al., 2004; Brown et al., 2005). Sensory feedback from new proprioceptors in the developing limbs may also participate in the respecification process, as found in the adult rat brain where the occurrence of novel sensory information can trigger neurotransmitter phenotype switching in postsynaptic central neurons (Dulcis et al., 2013).

In conclusion, the development of the limb neuromuscular apparatus and limb-based locomotion during *Xenopus* metamorphosis occurs in successive stages that involve a close functional relationship with the preexisting axial motor system until full autonomy is achieved (Combes et al., 2004) and, as shown here, is associated with changing underlying molecular patterning. Amongst the latter, neurotransmitter phenotype switching at the NMJ may enable limb motor axons to reach their appropriate muscle targets, which constitutes a novel and fundamental role for this process during motor innervation development (Spitzer, 2017) and adds to our understanding of NMJ development in general.

## MATERIALS AND METHODS

### Animals

Experiments were conducted on the South African clawed toad *X. laevis* obtained from the Xenopus Biology Resources Centre in France (University of Rennes 1; http://xenopus.univ-rennes1.fr/). Animals were maintained at 20–22°C in filtered water aquaria with a 12:12h light/dark cycle. Developmental stages were sorted according to external body criteria (Nieuwkoop and Faber, 1956), and experiments were performed on larvae from stage 49 to 57. All procedures were carried out in accordance with, and approved by, the local ethics committee (protocols #68-019 to HT and #2016011518042273 APAFIS #3612 to DLR).

### Motoneuron retrograde tracing

Procedures used for neuronal retrograde tracing were as described previously (Bougerol et al., 2015). Briefly, animals were anesthetized in a 0.05% MS-222 water solution and transferred into a Sylgard-lined Petri dish. In order to backfill MNs from their muscle targets, the skin covering the muscles of interest was dried before a tiny incision was made and crystals of fluorescent dextran amine dyes were applied intramuscularly (Forehand and Farel, 1982; van Mier et al., 1985; Roberts et al., 1999). In most cases, only hindlimb MNs were labeled with either 3 kD rhodamine (RDA) or 10 kD Alexa Fluor 647 (Thermo Fisher, Illkirch, France), except in the experiments illustrated in Fig. 1B, C where axial MNs were also labeled using 10 kD Alexa Fluor 488. Excess dye was washed out with cold Ringer solution (75 mM NaCl, 25 mM NaHCO_3_, 2 mM CaCl_2_, 2 mM KCl, 0.5 mM MgCl_2_, and 11 mM glucose, pH 7.4). After recovering from anesthesia, larvae were kept in a water tank for 24-48h to allow tracer migration into MN cell bodies and dendrites. In a series of experiments, the hindlimb buds of stages 51-52 larvae were injected and the animals were kept for several days in a separate aquarium until reaching metamorphic climax (Fig. 2E), in order to verify that early stage MNs were preserved through later development. Generally, such a labeling approach stains a large proportion of neurons that project processes within the bud muscles and thus potentially labels both MNs and sensory neurons (Forehand and Farel, 1982; van Mier et al., 1985; Roberts et al., 1999). However, since MNs have centrally located cell bodies, our retrograde tracings allowed the confident identification of MN somata only within the spinal cord.

### Immunofluorescence labeling

After MN retrograde labeling, spinal cords were dissected out and fixed in 4% paraformaldehyde (PFA) for 12h at 4°C. Preparations were incubated in a 20% [in phosphate-buffered saline (PBS) 0.1%] sucrose solution for 24h at 4°C, then embedded in a tissue-tek solution (VWR-Chemicals, Fontenay-sous-Bois, France) and frozen at −45°C in isopentane. 40 μm cross-sections were cut using a cryostat (CM 3050, Leica, Nanterre, France). Fluorescence immunohistochemistry was carried out on these spinal cross-sections using the same protocol as described previously (Bougerol et al., 2015). Briefly, after several rinsing steps and the blocking of non-specific sites (using a solution with PBS, Triton X-100 0.3%, bovine serum albumin 1%; Sigma, St. Quentin Fallavier, France) samples were incubated with the primary antibody for 48h at room temperature. After rinsing, cross-sections were incubated for 90 min at room temperature with a fluorescently labeled secondary antibody, and washed again before mounting in a homemade medium containing 74.9% glycerol, 25% Coon’s solution (0.1M NaCl and 0.01M diethyl-barbiturate sodium in PBS), 0.1% paraphenylenediamide. The primary antibodies used were goat anti-ChAT (1:100; Millipore), rabbit anti-VAChT (1:1000; Santa-Cruz) and mouse anti-Islet1/2 (1:250; Developmental Studies Hybridoma Banks (DHSB), University of Iowa, Iowa city, US). Secondary antibodies were donkey anti-goat and anti-rabbit or anti-goat and anti-mouse IgGs coupled to Alexa Fluor 488 and 568 (1:500; Life Technologies).

Whole-mount fluorescent immunohistochemistry was carried out on entire hindlimb buds from developmental stages 51 to 57 after overnight fixation in PFA 4%. The same labeling protocol was used as described above for spinal cross-sections. Alexa Fluor 488-conjugated α-bungarotoxin (10 μg/ml; Life Technologies) was used to label neuromuscular junctions. The primary antibody, mouse anti-neurofilament associated protein (3A10; 1:100; DSHB, Iowa city, Iowa, USA), was used to label nerve branches innervating the limb bud. The primary antibodies, rabbit anti-synaptophysin (1:500; abcam, Paris, France) and goat anti-NMDA subunit 1 receptor (NR1; 1:100; abcam) or a rabbit anti-NR2b (1:200; abcam), were used to label synapses and glutamate receptors, respectively. The bud preparations were then incubated with secondary antibodies donkey anti-mouse, anti-rabbit and anti-goat IgGs coupled to Alexa Fluor 488, 568 and 647 (1:500, Thermo Fisher). For microscope imaging limb buds were mounted on cavity slides in a homemade medium (see above).

### Image acquisition and fluorescence quantification

Whole-mount preparations and cross-sections labeled with fluorescent material were imaged using an Olympus FV1000 confocal microscope equipped with 488, 543 and 633 nm laser lines. Images were processed using Fiji and Photoshop softwares. Multi-image confocal stacks with 1 μm z-step intervals were generated using a 20x/0.75 oil objective and with 0.3 μm z-step intervals using a 60x/1.4 oil objective. Figure images were obtained by orthogonal projection from multi-image stacks with artificial fluorescent colors using the freeware Fiji.

ChAT, VAChT and Islet1/2 fluorescence quantification were performed on original images from 0.3 μm z-step stacks. Fluorescence intensity was measured automatically from 3 ROI in single planes with the same size, defining the slice background, and the axial and appendicular MNs fluorescence signals, respectively. The variation of fluorescence ((F-F_0_)/F_0_) was calculated for both axial and appendicular MNs ROI relative to background. This calculation was performed on 5 consecutive confocal planes per slice where the appendicular motor column was identified by retrograde labeling (usually 2 to 5 40-μm slices) and where background noise was maximal.

### In situ hybridization

Because of its tetraploid status, *X. laevis* possesses two genes for ChAT (Genbank accession numbers: XM_018225547 and XM_018239425 for ChAT1 and ChAT2, respectively) that exhibit about 87% nucleotide identity. This prompted us to generate two ChAT probes. To synthesize these probes, two PCR fragments of 529 (ChAT1) and 512 (ChAT2) bp were amplified from adult *X. laevis* brain and spinal cord RACE-ready cDNA (Bougerol et al., 2015), then subcloned into pGEM-T easy (Promega, Charbonnières, France). The pairs of primers used were: 5′-TTTGCTGCCAACCTTATCTCTG-3′ and 5′-ATGAAGTAACTGCAAAGCCCTG-3′ for ChAT1, and 5′-TCTGGAGTGCTGGATTACAA-3′ and 5′-ATGAAGTAACTGCAAAGCCCTG-3′, for ChAT2. Sense and antisense digoxigenin-labeled riboprobes were synthesized from the linearized plasmid with the RNA polymerases T7 or Sp6, using the RNA Labeling Kit (Roche Diagnostics, Mannheim, Germany).

The trunk region of tadpoles from stages 49 to 55 was dissected, fixed with 4% PFA in 0.1 M PBS overnight at 4°C and rinsed in 0.1 M PBS. Fixed samples were cryoprotected in 15% then 30% sucrose/PBS and embedded in Tissue-Tek (Sakura, Netherlands). Frontal sections (20 μM) of the trunks were cut at −20°C using a cryostat, collected on Superfrost Plus slides (O. Kindler, Freiburg, Germany), dried at room temperature for 24h and stored at −80°C until use.

The *in situ* hybridization protocol used in the present study was adapted from earlier studies (Buresi et al., 2012; Bougerol et al., 2015) and consisted of the following steps. Briefly, sections were rinsed 2×5 min with PBS at room temperature, 15 min in 5 times concentrated sodium chloride and sodium citrate solution (5X SSC), then were incubated for 2h in prehybridization buffer (50% formamide, 5X SSC, 50 μg/ml heparin, 5 mg/ml yeast RNA, 0.1% Tween) at 65°C. When prehybridization was complete, the prehybridization solution was removed and replaced with the same buffer containing a mix of the two heat-denaturated digoxigenin-labeled ChAT1 and ChAT2 riboprobes. Hybridization was carried out overnight at 65°C. Sections were rinsed 3×30 min in 2X SSC at 65°C then 1h in 0.1 X SSC at 65°C. Two final washes were performed for 5 min in MABT (maleic acid 100 mM, pH 7.2, NaCl 150 mM, Tween 0.1%) at room temperature. Sections were transferred to a blocking solution [5% blocking reagent (Roche), 5% normal goat serum in MABT] and incubated at room temperature for 1h, before addition of the alkaline phosphatase coupled anti-digoxigenin antibody (1/4,000) for overnight storage at 4°C. Sections were again washed 3×10 min in MABT and 2×5 min in PBS at room temperature, then incubated 10 min in staining buffer (100 mM Tris-HCl, pH 9.5, 50 mM MgCl_2_, 100 mM NaCl, and 0.1% Tween) and transferred to BM Purple (Roche) for colorimetric detection. Finally, they were washed twice in PBS to stop the reaction, and then mounted on gelatin-coated slides in Mowiol. Sections were imaged using a Leica DM5500 B microscope connected to LAS V4.1 software. The specificity of the hybridization procedure was verified by incubating sections with the sense riboprobes.

### Patch-clamp recording of appendicular motoneurons

Retrograde labeling of appendicular MNs was performed using 3kD RDA dextran on stages 51, 52 larvae. The day after, patch-clamp electrophysiological recordings of labeled MNs were made on isolated brainstem-spinal cord *in vitro* preparations. After anesthesia in 0.05% MS-222, the brainstem and spinal cord, including spinal segmental ventral roots, were dissected out in cold oxygenated (95% O_2_, 5% CO_2_) Ringer solution. The preparation was then placed in a recording chamber and continuously superfused with oxygenated Ringer solution (~ 2.5 mL, ~17°C, rate of ~2 mL/min). Spontaneous fictive locomotor episodes were recorded from a caudal ventral root (between segments 12 and 15; Fig. 3B_1_) using a borosilicate glass suction electrode (tip diameter, 100 nm; Clark GC 120F; Harvard Apparatus) filled with Ringer solution. The recorded signal was amplified (A-M system), rectified and integrated (time constant 100 ms; Neurolog System). RDA-positive appendicular MNs were identified with a standard epifluorescent illumination system (Cy3 filter) within the whole-mount spinal cord (dorsal-side opened) and subsequently visualized using a differential interference contrast microscope with an infrared video camera to facilitate the patch electrode trajectory (Fig. 3B). Using an Axoclamp 2A amplifier (Molecular Devices, Berkshire, UK), whole-cell patch-clamp recordings were made with a borosilicate glass electrode (pipette resistance, 5-6 MΩ; Clark GC 150TF; Harvard Apparatus) filled with a solution containing 100 mM K-gluconate, 10 mM EGTA, 2 mM MgCl_2_, 3 mM Na_2_ATP, 0.5 mM NaGTP, 10 mM HEPES, pH 7.3. All electrophysiological signals were computer-stored using a digitizer interface (Digidata 1440; Pclamp10 software; Molecular Devices) and analyzed offline with Clampfit software (Molecular Devices).

### Axial and hindlimb EMG recordings in semi-intact preparations

Semi-intact preparations from stages 52 to 57 larvae were used to simultaneously record EMG activity from axial myotomes and hindlimb bud muscles (Fig. 4A). Brainstem-spinal cord preparations were dissected out in the same way as for patch-clamp recording, but tail myotomes (7-10) and hindlimb buds were left attached to the spinal cord. Semi-intact preparations were fixed in a Sylgard-lined recording chamber, continuously superfused with oxygenated Ringer solution (1.3–2.1 mL/min) and maintained at 18±0.1°C with a Peltier cooling system. In some experiments, *d*-tubocurarine (30 μM; Sigma) was exogenously applied to block nicotinic receptor type cholinergic synapses. Fictive locomotion sequences were generated either spontaneously or triggered by electrical stimulation of the caudal region of the brainstem (Grass stimulator S88), and the spinal swimming pattern was monitored from a caudal ventral root (between segments 12 and 15) with a suction electrode as described above. EMG activities in rostral myotomes (7-10) and hindlimb buds were recorded simultaneously using pairs of 50 μm insulated wire electrodes connected to a differential AC amplifier (A-M System). Both nerve and EMG activities were digitized at 10 kHz (CED 1401, Cambridge Electronic Design, UK), and displayed and stored on computer for offline analysis with Spike2 software (CED).

Discharge rates in individual nerve and EMG recordings were measured by setting an amplitude threshold to count all impulses in such multi-unit recordings. Firing rates (in spikes/s) were averaged over 10-20 locomotor cycles. Cycle period was taken as the interval between the onsets of two consecutive ventral root bursts. These consecutive burst onsets were used as a trigger for averaging the discharge rates of each EMG channel over cycle duration.

### EMG recordings in isolated nerve-limb bud preparations

Isolated hindlimb-bud preparations with the sciatic and crural nerves still attached (Fig. 5C) were used to record the EMG activity evoked by electrical stimulation of either appendicular nerve branch at stages 52 to 57. Under MS-222 anesthesia, appendicular nerves were disconnected from the spinal cord and separated from tail myotomes, taking care not to detach them from the rest of the bud. The bud and attached nerves were fixed with small pins in a Sylgard-lined recording chamber and superfused with oxygenated Ringer solution. In some experiments, 30 μM *d*-tubocurarine or a cocktail of 20 μM CNQX + 10 μM AP5 was added to the perfusion solution to block nicotinic or glutamatergic receptors, respectively (all drugs came from Sigma). A small incision was made at the distal extremity of the bud to allowing insertion of the EMG electrodes. Either limb nerve branch was stimulated with a glass suction electrode connected to a Grass stimulator S88 through a photoelectric stimulus isolation unit (PSIU 6; Grass Instruments). Note that stimulating either branch provided similar results, and no distinction was made in this report. Single pulses (70-300 μA; 10 μs) were delivered every 100s, and polarity inversion tests were performed in order to distinguish the stimulation artifact from the muscle response. EMG signals were amplified, integrated (time constant 10 ms) with Spike2 software and stored as described above. EMG responses were measured as the area under the integrated recording traces.

### Statistics

After signal processing in Spike2, electrophysiological data were analyzed using Prism5 (GraphPad, USA) and OriginPro8 (OriginLab Corporation, USA). Data are shown as means and standard errors of the mean (+SEM), unless stated otherwise. For integrated EMG signals, differences of means were tested using the unpaired two-tailed Mann–Whitney U-test.

## Acknowledgements

We are grateful to Gilles Courtand (UMR 5287) for technical assistance in image analysis, and Feng Quan (UMR 7221) and Boudjema Imarazene (UMR 7208) for technical assistance with *in situ* hybridization. This work was supported by grants from the *Centre National de la Recherche Scientifique* and by the *Actions Thématiques du Muséum*.

